# Cytoskeletal disassembly by optogenetic control of RhoA signaling termination

**DOI:** 10.64898/2026.09.03.749286

**Authors:** Erin E. Berlew, Jude Barakat, Paula Camacho Sierra, Joel D. Boerckel

## Abstract

Cellular morphodynamics require adaptive cytoskeletal remodeling, mediated by precisely coordinated activation and termination of RhoA GTPase signaling. RhoA activation is well-studied, but the kinetics and molecular basis of signaling termination remain poorly understood. We engineered an optogenetic toolbox on the single-component BcLOV4 platform for bidirectional control of RhoA activation (opto-GEF11) and termination (opto-DLC1). Prior studies have inferred GTPase inactivation kinetics indirectly, by tracking passive recovery from an activated state, but whether this reflects the kinetics of forward signaling termination remains unclear. Using opto-DLC1, we show that direct RhoA termination is an order of magnitude faster than passive disactivation, despite comparable signaling amplitude. Mechanistically, opto-DLC1 triggered rapid actin disassembly through cofilin disinhibition but drove YAP nuclear efflux at half the rate of opto-GEF11-induced nuclear influx. Together, these findings introduce a platform technology for controlling protein signaling termination, resolve RhoA activation and termination kinetics with sub-second precision, and reveal a mechanistic asymmetry between signaling activation and termination that enables cytoskeletal homeostasis.

## INTRODUCTION

Cellular morphodynamics mediate tissue development, repair and homeostasis, and require a dynamic actin cytoskeleton that provides structural integrity but also remains responsively adaptative to biochemical and mechanical stimuli^1,2^. Actin cytoskeletal dynamics are mediated by the Rho (Ras homology) family of small GTPases, which signal at the plasma membrane and conduct an array of downstream events to generate morphodynamic forces, polymerize and remodel the actin cytoskeleton, and transduce mechanical cues into biological signals^3-5^. GTPases toggle between active (GTP-bound) and inactive (GDP-bound) states through activating GEFs (Guanine exchange factors) and inactivating GAPs (Guanine accelerating proteins)^6^ (**Fig. 1A**). Recent work on the Rho family GTPases has emphasized the diversity of GEF/GAP interactions with the GTPases^7,8^, their subcellular distributions^9^, and associated cytoskeletal outcomes^10^. The emergence of optogenetic tools that enable precise spatiotemporal control of signaling activation has provided important insights into the kinetics and spatial patterning of GTPase signaling in cell motility, mechanotransduction, and cytokinesis^11-15^; however, the biology and dynamics of Rho signaling termination remain poorly understood. Here, we engineer a suite of optogenetic tools for dynamic control of RhoA signaling termination, resolve termination kinetics with sub-second precision, and define the molecular basis of downstream cytoskeletal homeostasis.

**Figure 1.**
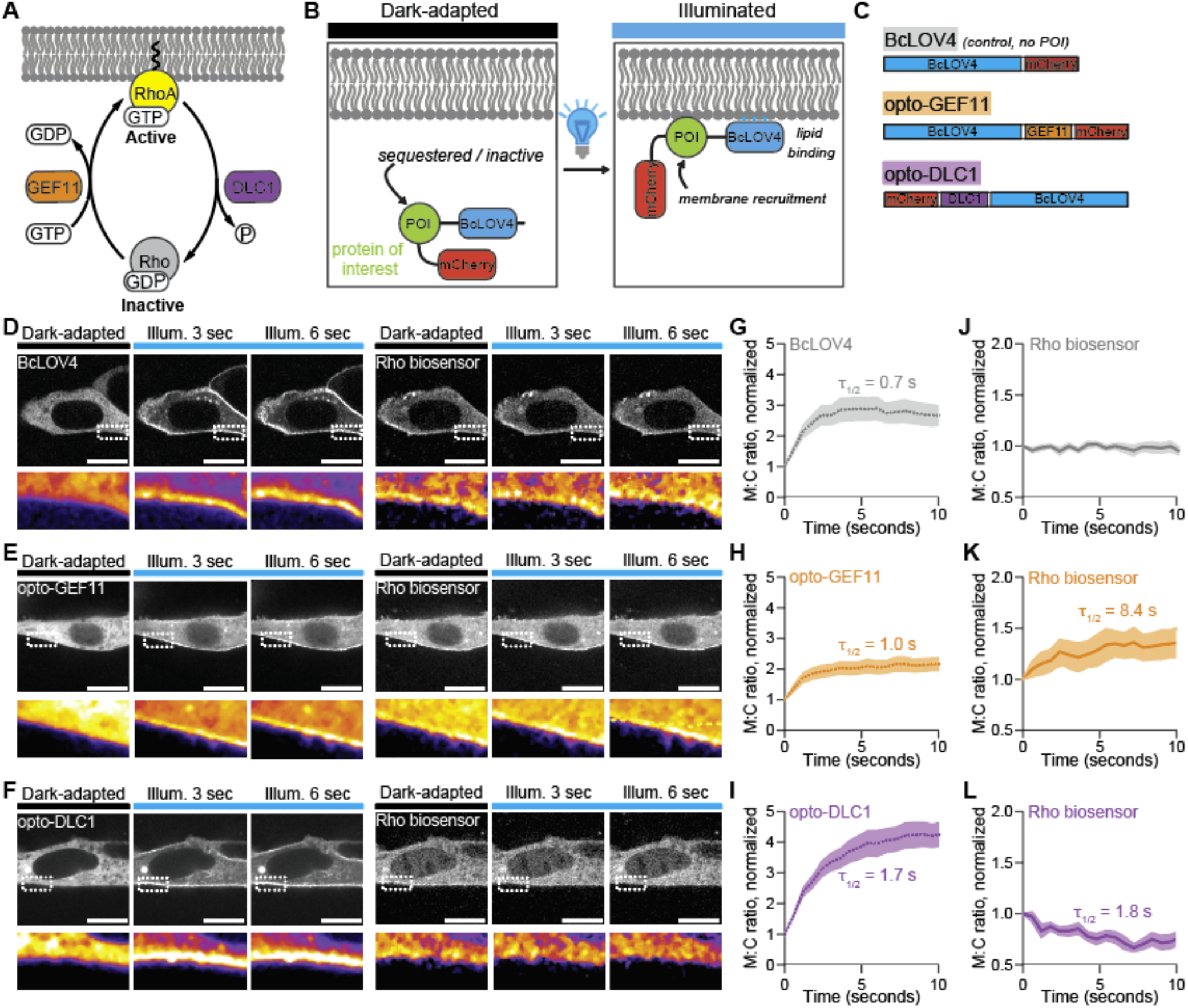
Optogenetic control of RhoA activation and termination. **A**) Schematic representation of RhoA GTPase signaling at the plasma membrane. **B**) Optogenetic tool engineering strategy for light-induced membrane recruitment of BcLOV4-fused proteins of interest (POI). **C**) Block diagram of optogenetic tools. BcLOV4, mCherry, and GEF/GAP domains were separated by flexible (GGGS)_2_ linkers to create BcLOV4 control, opto-GEF11, and opto-DLC1 tools. **D-F**) Representative images of BcLOV4, opto-GEF11, and opto-DLC1, visualized by eGFP-tagged tool (left) and dTomato-tagged Rho biosensor (right). Scale = 5 µm. Inset shows boxed region in perceptually uniform color palette. **G-I**) Quantification of BcLOV4, opto-GEF11, and opto-DLC1 tool membrane-to-cytosol fluorescence ratio. **J-L**) Quantification of Rho biosensor membrane-to-cytosol fluorescence ratio following BcLOV4, opto-GEF11, and opto-DLC1 stimulation. Mean ± S.E.M, N = 7-15 cells per condition. Time constants calculated from one-phase association exponential fit.

The Rho family GTPase RhoA mediates cytoskeletal contractility and stress fiber formation^3,5,16,17^. RhoA signaling generates cytoskeletal tension via Rho-associated protein kinase (ROCK)-mediated myosin motor phosphorylation and activates formins for ROCK-independent actin polymerization, resulting in dynamic stress fiber formation^5,18^. These actions are critical for diverse cellular functions, including cell adhesion, motility and cytokinesis; however, persistent or excessive activation of RhoA signaling can cause cytoskeletal arrest. For example, disruption of the transcriptional programs that produce RhoA-inactivating GAPs causes over-activation of RhoA signaling, impaired cytoskeletal and adhesion remodeling, and progressive motility arrest in both cells and embryos^19-22^. Functional cytoskeletal homeostasis therefore requires feedback regulation of RhoA signaling; thus, dynamic systems under feedback control must be studied in the time-domain and cannot be understood by examining equilibrium states alone.

However, tools and techniques with which to study the biology of signaling termination with spatiotemporal resolution are limited. Global manipulations like genetic protein depletion/deletion or overexpression are not designed to reveal spatiotemporal detail^21-24^. Pharmacological inhibitors that target components of Rho signaling have been impactful, both mechanistically and therapeutically^25^; however, these tools lack spatial control and their temporal kinetics are rate-limited by transport time scales that are longer than theoretical time scales of local signaling control. Further, available inhibitors and signaling agonists target signaling cascade events up- or down-stream of RhoA, but do not interrogate the biology of GAP-mediated Rho-GTP hydrolysis. For example, inhibitors like Rhosin prevent RhoA activation rather than terminating signaling specificity^26^, and commonly-used compounds like Y27632 interfere with the kinase activity of downstream ROCK rather than modifying GTPase signaling *per se*^27,28^. To date, experimental measurements of RhoA signaling termination kinetics in mammalian cells have not been provided.

Optogenetic tools provide laser-pulse precision for spatiotemporal dissection of GTPase signaling by mimicking native protein-protein interactions that activate and terminate signaling events. To date, the optogenetics field has focused predominantly on the kinetics of signaling activation^11,12,15,29-35^, but termination remains poorly understood. Current methods to study signaling inactivation dynamics in mammalian cells are indirect. For example, activating tools have been used to infer the kinetics of “disactivation”, i.e., passive recovery from an activated state^12,13^, but the kinetics and consequences of direct signaling termination have not been explored.

Here, we introduce opto-DLC1, a blue light-inducible RhoA signaling termination tool. DLC1 (Deleted in liver cancer 1, also known as ARHGAP7) is a RhoA GTPase that regulates actin stress fiber formation and mechanotransductive cytoskeletal contractility^36,37^. We characterize opto-DLC1, in experimental contrast with our previously reported RhoA activation tool, opto-GEF11^31^, and quantify the temporal kinetics and cytoskeletal and mechanotransductive consequences of both RhoA activation and termination. Further, we show that RhoA termination kinetics are an order-of-magnitude faster than RhoA disactivation kinetics, despite similar signaling magnitude shifts, and demonstrate mechanistically that RhoA termination drives rapid actin depolymerization via cofilin disinhibition.

## RESULTS

### Engineering optogenetic tools for bi-directional RhoA activation and termination

BcLOV4 is a fungal-derived photosensor protein that exhibits natural blue light-induced lipid binding^32,38^ and clustering^39^. We previously established the BcLOV4 optogenetic platform, in which blue light stimulation triggers BcLOV4/protein-of-interest (POI) fusion protein translocation from the cytosol to the plasma membrane inner leaflet ^31,40,41^ (**Fig. 1B**). Here, we engineered opto-DLC1 for inducible RhoA signaling termination by fusing the catalytic GAP domain of DLC1/ARHGAP7 with BcLOV4 and an mCherry visualization tag (mCherry-DLC1-BcLOV4-3xFLAG) (**Suppl. Fig. 1**). This platform technology is generalizable for ARHGAP signaling, which we demonstrate with opto-GAP1 (**Suppl. Fig. 2**), an alternate RhoA-inactivating GAP^42,43^. We contrast opto-DLC1-mediated termination of endogenous RhoA GTPase signaling with opto-GEF11 (**Suppl. Fig. 2**), which activates endogenous RhoA via BcLOV4-mediated membrane recruitment of the catalytic GEF domain of ARHGEF11.

**Figure 2.**
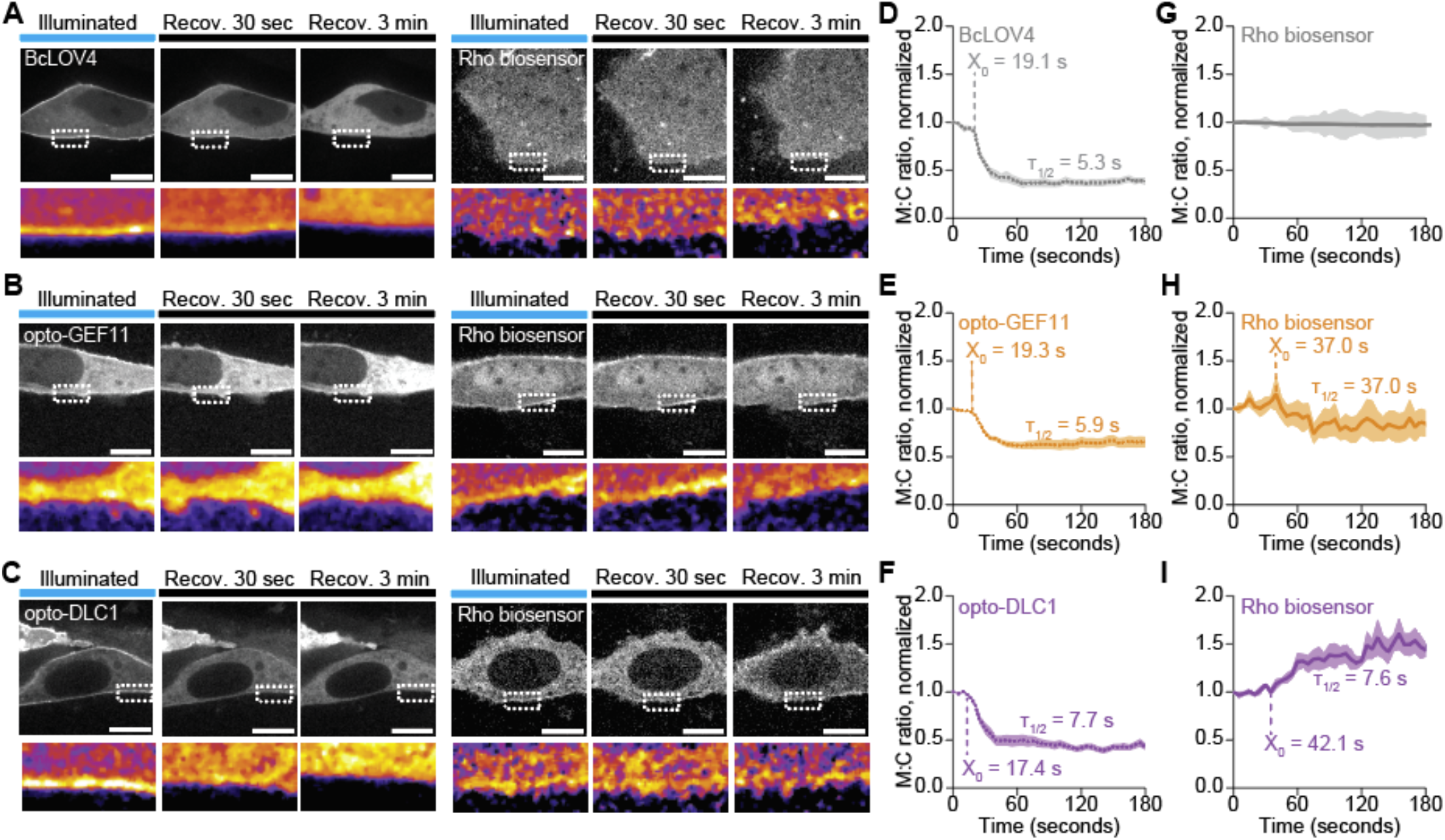
Kinetics of passive recovery from activated and terminated states. **A-C**) Representative images of BcLOV4, opto-GEF11, and opto-DLC1, visualized by eGFP-tagged tool (left) and dTomato-tagged Rho biosensor (right), during dark-state recovery from blue-light stimulation. Scale = 5 µm. Inset shows boxed region in perceptually uniform color palette. **D-F**) Quantification of BcLOV4, opto-GEF11, and opto-DLC1 tool membrane-to-cytosol fluorescence ratio during dark-state recovery from blue-light stimulation. **G-I**) Quantification of Rho biosensor membrane-to-cytosol fluorescence ratio following cessation of BcLOV4, opto-GEF11, and opto-DLC1 stimulation, respectively. Mean ± S.E.M, N = 3-15 cells per condition. Time constants calculated from delay plateau followed by one-phase dissociation exponential fit. X_o_ = length of delay plateau.

### Kinetics of RhoA activation and termination

To measure the kinetics of GEF-mediated RhoA activation and GAP-mediated RhoA termination, we co-expressed a localization-based Rho biosensor (dTomato-2xrGBD), which preferentially binds active GTP-bound RhoA^44^, with opto-GEF1, opto-DLC1, and BcLOV4 as control (**Fig. 1D-F**). We measured membrane-to-cytosol fluorescence for both biosensor and tool during pulsatile blue light stimulation and quantified dynamics by one-phase decay curve-fit to calculate half-time to equilibrium (τ_1/2_). Blue light stimulation induced similarly rapid membrane association for BcLOV4 control (τ_1/2_ = 0.7 sec), opto-GEF11 (τ_1/2_ = 1.0 sec) and opto-DLC1 (τ_1/2_ = 1.7 sec) (**Fig. 1G-I**). BcLOV4 control stimulation did not alter plasma membrane Rho signaling, measured by membrane/cytosolic Rho biosensor localization (**Fig. 1J**). Opto-GEF11 activated Rho signaling, with τ_1/2_ = 8.4 sec (**Fig. 1K**), while opto-DLC1 terminated Rho signaling, with τ_1/2_ = 1.8 seconds (**Fig. 1L**).

### Kinetics of passive recovery from activated and terminated states

Prior studies have inferred the kinetics of GTPase signaling inactivation by tracking passive recovery from an activated state^12,13^, but whether this reflects the kinetics of forward signaling termination remains unknown. Therefore, we measured the kinetics of tool dissociation and RhoA signaling recovery to basal state after one minute of pulsatile stimulation for BcLOV4 control, opto-GEF11, and opto-DLC1 (**Fig. 2**). We refer to passive recovery from an activated RhoA signaling state as “*disactivation*” and passive recovery from a terminated RhoA signaling state as “*distermination*”. Tool localization kinetics were measured using mCherry variants of BcLOV4, opto-GEF11, and opto-DLC1, and Rho signaling kinetics were measured using dTomato-tagged biosensor with co-expressed eGFP variants of the tools to enable live imaging after blue light cessation (**Fig. 2A-C**). For all tools, disassociation from the membrane exhibited a time delay and was modeled by a plateau followed by exponential decay (BcLOV4: delay = 19.1 sec, τ_1/2_ = 5.3 sec; opto-GEF11: delay = 19.3 sec, τ_1/2_ = 5.9 sec; opto-DLC1: delay = 17.4 sec, τ_1/2_ = 7.7 sec) (**Fig. 2D-F**). For Rho biosensor localization, we measured no change for BcLOV4 controls, as expected (**Fig. 2G**). RhoA signaling disactivation, or recovery from an opto-GEF11 activated state, exhibited a decay half-time of τ_1/2_ = 37.0 sec, after a delay of 37.0 seconds (**Fig. 2H**). RhoA signaling distermination, or recovery from an opto-DLC1 terminated state, exhibited a half-time of τ_1/2_ = 7.6 sec, after a delay of 42.1 seconds (**Fig. 2I**).

### Optogenetic RhoA termination and activation out-perform passive recovery in kinetics and potency, respectively

To distinguish direct RhoA signaling termination vs. passive disactivation and direct RhoA activation vs. passive distermination (**Fig. 3A**), we quantitatively compared nonlinear regression kinetics (half-time) and magnitudes (plateau) by Extra Sum-of-Squares F-tests. Opto-DLC1 termination was significantly faster (**Fig. 3B**), but equivalent in plateau amplitude (**Fig. 3C**), compared to passive disactivation from an opto-GEF11 activated state. In contrast, opto-GEF11 activation was equivalent in kinetics (**Fig. 3B**), but greater in plateau amplitude (**Fig. 3C**), compared to passive activation from an opto-DLC1 terminated state. *Nota bene*: for time-scale comparisons, we did not include the time delay in passive recovery initiation, as this is likely driven by tool-membrane dissociation kinetics^32,38^.

**Figure 3.**
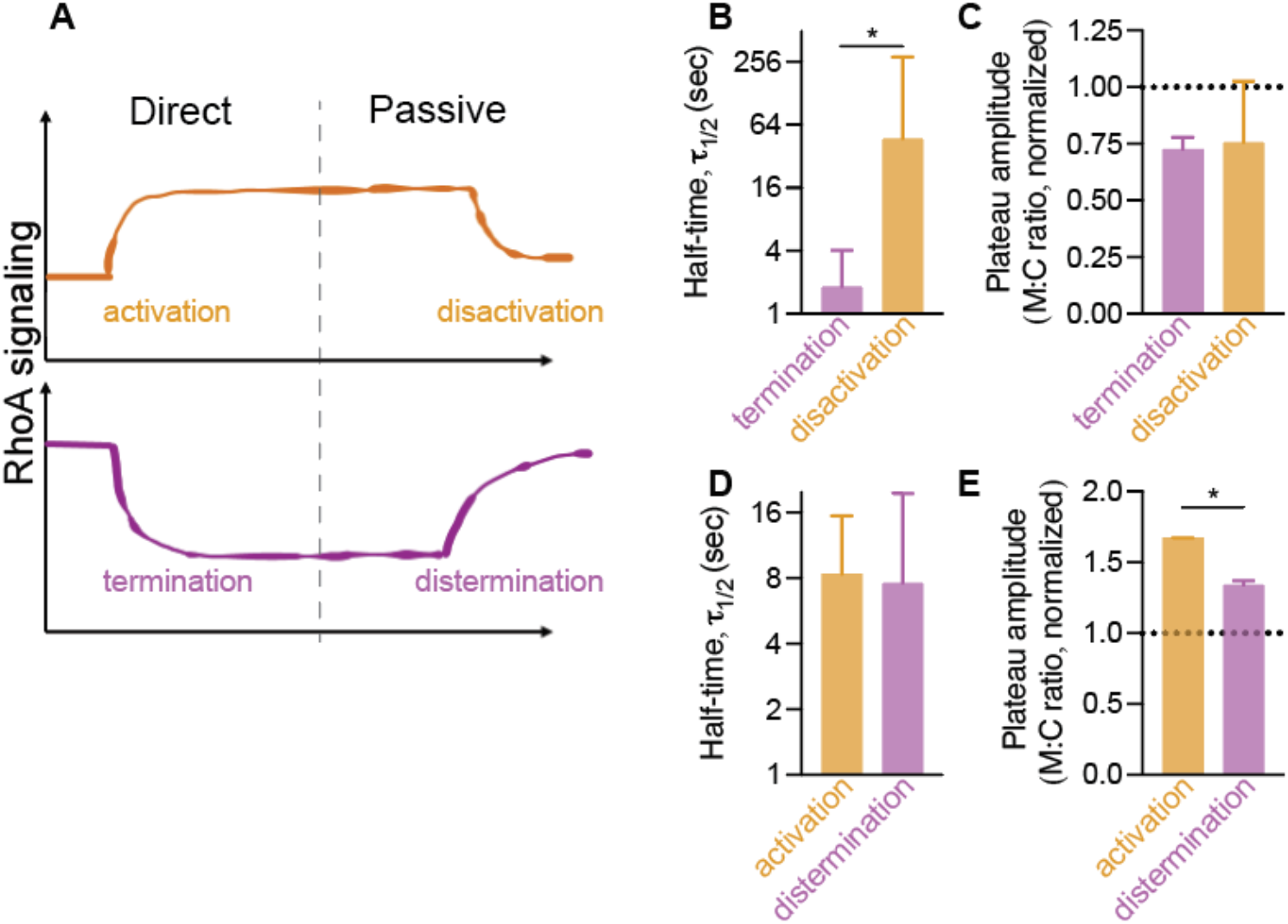
Passive recovery to basal state is not equivalent to activation or termination. **A**) Schematic illustration of direct activation dynamics and passive disactivation dynamics enabled by opto-GEF11, and direct termination dynamics and passive distermination dynamics enabled by opto-DLC1. **B,C**) Comparison of half-time and plateau amplitude, normalized to initial state (dotted line), for termination vs. disactivation. **D,E**) Comparison of half-time and plateau amplitude, normalized to initial state (dotted line), for activation vs. distermination. * indicates p < 0.05, Extra Sum-of-Squares F-test.

### Bidirectional optogenetic control of mechanotransductive cytoskeletal contractility and actin dynamics

We previously showed that optogenetic RhoA activation, either by membrane recruitment of the RhoA GTPase (opto-RhoA) or of ARHGEF11 (opto-GEF11), induced mechanotransductive cytoskeletal contractility, measured by dynamic nuclear localization and transcriptional activity of the mechanoresponsive transcriptional regulator, YAP (Yes-associated protein)^31^. Here, we mapped YAP nucleocytoplasmic localization in response to both RhoA activation and termination (**Fig. 4**). We quantified the nuclear:cytoplasmic ratio of eGFP-tagged YAP, co-expressed with either opto-GEF11 or opto-DLC1, over a time course of 30 minutes of pulsatile blue light stimulation. Opto-GEF11 drove YAP nuclear import (τ_1/2_ = 12.6 min), after tool recruitment to the plasma membrane (**Fig. 4A,C**). Conversely, opto-DLC1 induced YAP nuclear efflux (τ_1/2_ = 22.5 min), after a delay plateau of 8.6 min (**Fig. 4B,C**). Together, these data indicate dynamic regulation of cytoskeletal contractility.

**Figure 4.**
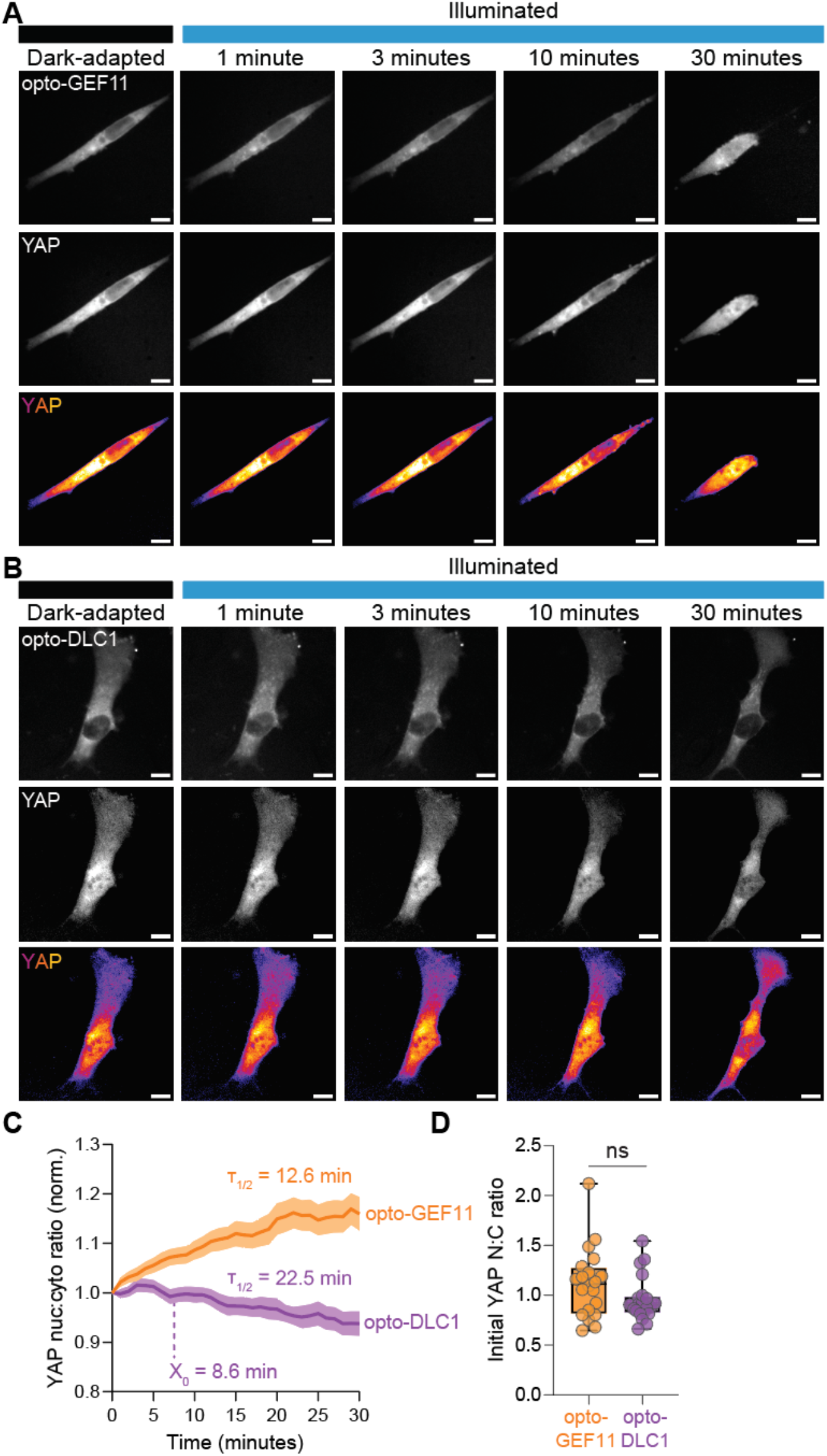
RhoA activation and termination dynamically control mechanotransductive cell contractility. **A**) Representative images of opto-GEF11 tool and eGFP-tagged YAP during a 30-minute period of pulsatile blue light stimulation. **B**) Representative images of opto-DLC1 tool and eGFP-tagged YAP during a 30-minute period of pulsatile blue light stimulation. Scale bars = 10 µm. **C**) Nuclear to cytosolic YAP intensity ratio, normalized to dark-adapted state. Mean ± S.E.M, N = 20-21 cells per condition. Time constants calculated by one-phase association (opto-GEF11) and plateau followed by one-phase dissociation (opto-DLC1) exponential fit. X_o_ = length of delay plateau.

Next, we visualized fluorescently tagged LifeAct to measure the kinetics of RhoA-regulated cytoskeletal remodeling (**Fig. 5**). Opto-GEF11 recruitment rapidly increased stress fiber formation and F-actin abundance (τ_1/2_ = 2.0 min) (**Fig. 5A,C**), followed by significant cellular contraction (τ_1/2_ = 6.3 min) (**Suppl. Fig. 3**). Opto-DLC1 recruitment induced cytoskeletal rearrangement and stress fiber disassembly, consistent with decreased ROCK-myosin contractility, but also caused a rapid and robust decrease in total F-actin (τ_1/2_ = 3.4 min) (**Fig. 5B,C**), suggesting that DLC1-mediated RhoA signaling termination also plays an active role in actin depolymerization.

**Figure 5.**
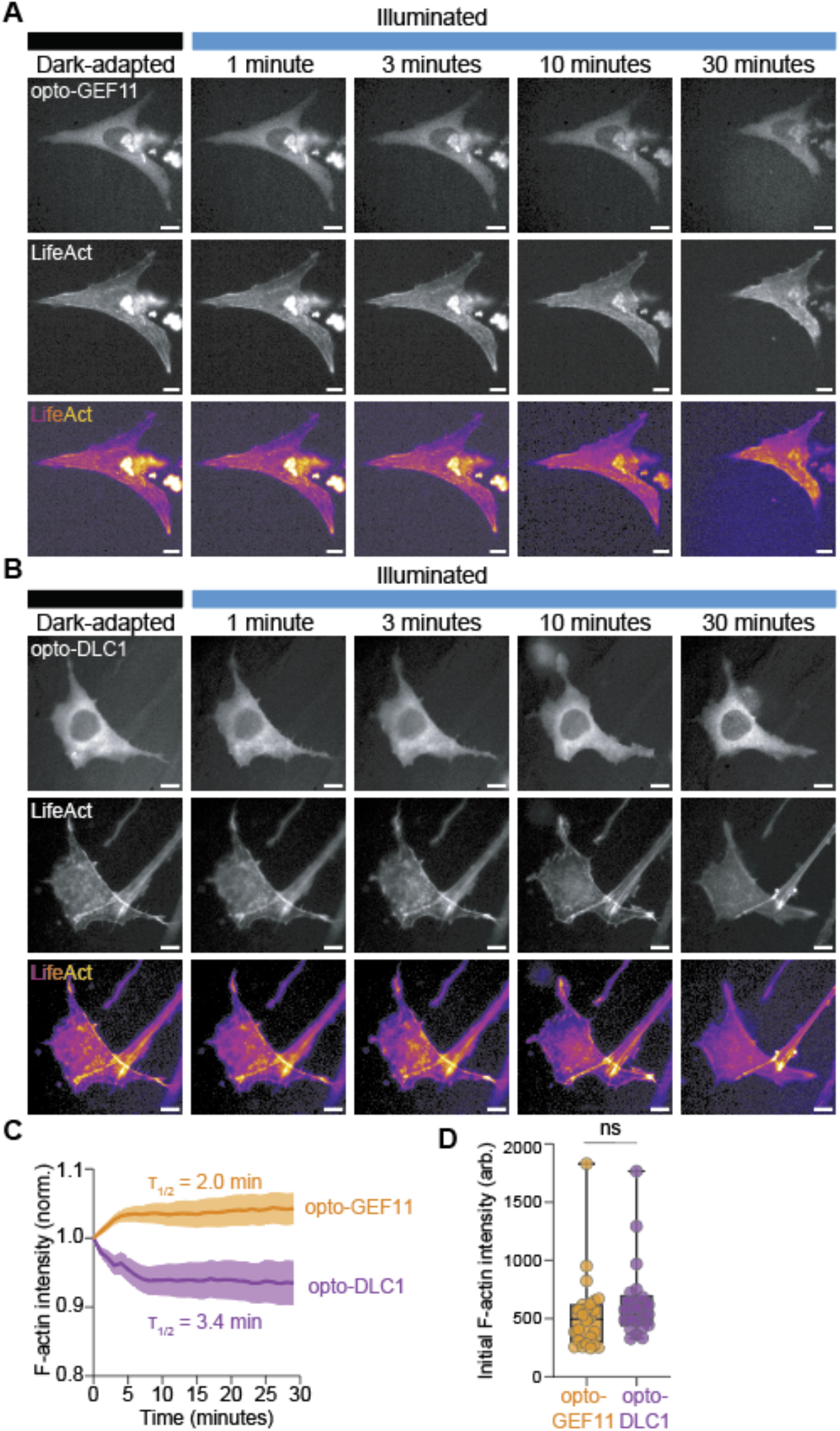
RhoA activation and termination dynamically control actin remodeling. **A**) Representative images of opto-GEF11 tool and miRFP703-tagged LifeAct during a 30-minute period of pulsatile blue light stimulation. **B**) Representative images of opto-DLC1 tool and miRFP703-tagged LifeAct during a 30-minute period of pulsatile blue light stimulation. Scale bars = 10 µm. **C**) F-actin intensity, normalized to dark-adapted state. **D**) Initial dark-state F-actin intensity. Mean ± S.E.M, N = 24, 26 cells per condition. Time constants calculated by one-phase association (opto-GEF11) and dissociation (opto-DLC1) exponential fit.

### Cofilin disinhibition induces rapid actin depolymerization following DLC1 recruitment

We next sought to identify the molecular mechanism of optogenetic DLC1-mediated actin depolymerization. We treated opto-DLC1-stimulated cells with jasplakinolide (Jasp), an inhibitor of actin depolymerization^45^, and found that Jasp-mediated filament stabilization prevented opto-DLC1-induced actin de-tensioning, demonstrating that the observed decrease in F-actin is due to disassembly of existing filaments rather than a lack of accumulation of new filaments (**Suppl. Fig. 4**). Next, we treated cells with D3, an inhibitor of Slingshot (SSH), a phosphatase that removes an inhibitory phosphate on the actin-severing protein cofilin^46^. While opto-DLC1 stimulation decreased F-actin abundance in DMSO-treated control cells, DLC1-induced actin depolymerization was abrogated by D3 treatment (**Fig. 6**). Together, these data identify DLC1 as a dynamic regulator of cellular morphodynamics that actively promotes actin disassembly via SSH disinhibition of cofilin.

**Figure 6.**
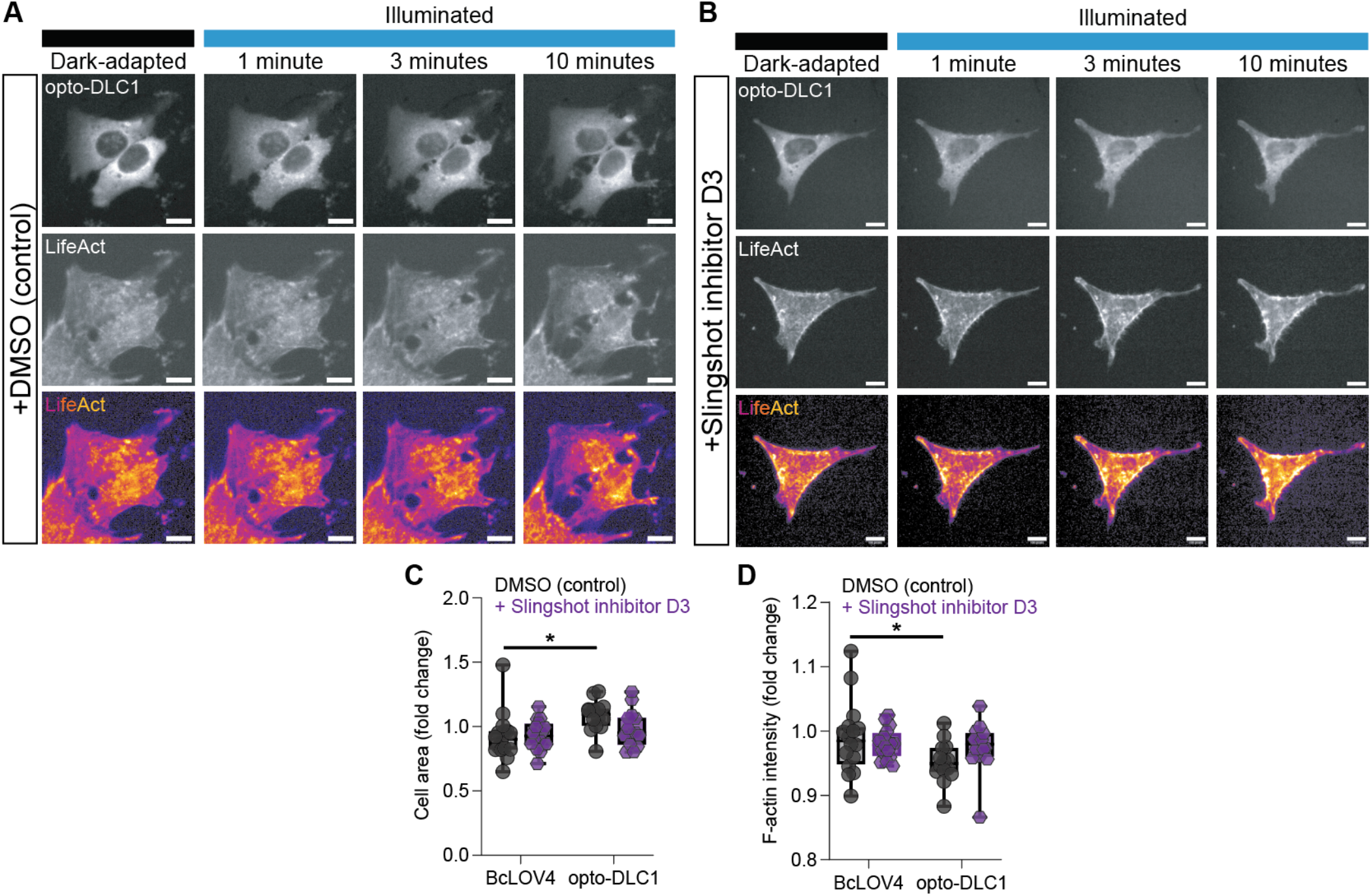
RhoA termination by Opto-DLC1 induces rapid actin disassembly via Slingshot-mediated cofilin disinhibition. **A,B**) Representative images of opto-DLC1 tool (top) and miRFP703-tagged LifeAct (bottom) during a 10-minute period of pulsatile blue light stimulation in the presence of DMSO (**A**) or slingshot inhibitor D3 (**B**). Scale = 10 µm. **C,D**). Quantification of cell area (**C**) and F-actin intensity (**D**). N = 14, 16 cells per condition. (*) p < 0.05 by two-way ANOVA with post-hoc comparisons by Šidák correction.

## DISCUSSION

Our understanding of the kinetics and biology of Rho GTPase signaling termination is limited by the spatiotemporal precision of tools and techniques currently available. Here, we engineer a bidirectional optogenetic toolbox for dynamic control of RhoA signaling activation and termination. We use these tools, together with live morphometry of downstream signaling and cytoskeletal effectors, to resolve RhoA activation and termination kinetics with sub-second precision. We present two approaches to studying bidirectional signaling. Specifically, we distinguish direct signaling termination from passive signaling disactivation and distinguish direct signaling activation from passive signaling distermination. Our data demonstrate that direct RhoA termination kinetics are an order-of-magnitude faster than passive disactivation kinetics, despite featuring a similar magnitude of decreased signaling at equilibrium. In contrast, RhoA activation features similar kinetics, but greater activation magnitude, compared to passive distermination. Functionally, opto-DLC1 induces cytoskeletal relaxation, YAP nuclear efflux, and notably rapid actin depolymerization, which could be abrogated by blocking actin severing by actin binding protein, cofilin. Together, these data define the kinetics of RhoA signaling activation and termination kinetics in mammalian cells and define a molecular basis for downstream cytoskeletal homeostasis.

Here, we used BcLOV4 to engineer a generalizable single-component optogenetic platform for Rho GTPase signaling termination. We demonstrate GAP modularity, introducing both opto-DLC1 and opto-GAP1 for light-inducible RhoA termination. To our knowledge, these represent the first optogenetic tools for GTPase inactivation in mammalian cells, though other systems exist for use in *Drosophila* (opto-GAP1)^47^ and for heterotrimeric GTPases (opto-RGS)^48^. We demonstrate the viability of recruiting a GAP domain to the plasma membrane to terminate membrane-associated GTPase signaling. As single component tools, which rely on light-induced structural rearrangement of the BcLOV4 amphipathic helix, resulting in reversible electrostatic binding to the inner leaflet of the plasma membrane, implementation requires expression of only a single plasmid^31,32^. This circumvents the necessity of stoichiometry-tuned plasmid co-expression, as for other multi-component tools such as iLID or CRY2-CIBN. These and other optogenetic platforms have been transformative for our current understanding of the spatiotemporal biology of RhoA signaling activation; however, signaling termination remains largely unexplored.

Using an optogenetic approach to directly recruit a RhoA-GTP hydrolyzing enzyme to the membrane, we were able to probe the kinetics and biology of RhoA signaling termination with previously inaccessible precision. In our prior study, using an opto-RhoA tool, also constructed on the BcLOV4 platform, we observed that RhoA accumulation at the membrane persists well after control-BcLOV4 dissociation^31^. We posited that signaling inactivation kinetics are likely governed by RhoA signaling dynamics rather than by comparatively rapid tool dissociation, but we lacked the ability to quantify the signaling and downstream effector dynamics. Our prior findings also raised the question of whether the kinetics of RhoA signaling termination by enzymatic GTP hydrolysis are distinct from those of passive recovery. Inspired by prior studies that measured downstream effector recovery kinetics after cessation of signaling activation by an opto-GEF, we designed tools and experiments that allow direct comparison.

We distinguish between direct RhoA signaling termination and passive disactivation, and between direct signaling activation and passive distermination. We observed significantly faster kinetics for opto-DLC1-mediated RhoA termination than for passive disactivation after cessation of opto-GEF11 (τ_1/2_ = 1.8 sec vs. 37 sec), despite similar plateau amplitude at equilibrium. As the basal equilibrium state is bounded by the physiological setpoint determined by cell type, mechanical environment, biochemical milieu, etc., we reason that the opto-DLC1 tool acts within the physiologic range of endogenous RhoA signaling. In contrast, opto-GEF11-mediated RhoA activation featured similar kinetics to passive distermination, suggesting that endogenous recovery to steady state involves similar GEF-mediated mechanisms. However, opto-GEF11-mediated RhoA activation exhibited a larger plateau amplitude than opto-DLC1-mediated RhoA distermination, suggesting that opto-GEF11 can induce supraphysiologic RhoA activity. We note that we did not take the X_0_ delay into account in our assessment of disactivation and distermination kinetics as this delay corresponded with the temporal kinetics of tool dissociation, but this could also represent meaningful interactions between activated RhoA or other effector complexes^49^. This additional variable further highlights the complexity inherent in indirect inference of termination kinetics and underscore the importance of engineering tools specifically to study signal termination. Using tools engineered on the same platform further facilitated direct comparison of kinetics: each tool could be expressed via a single plasmid and stimulated using the same intensities and duty ratios of blue light.

Our optogenetic approach further enabled previously inaccessible insight into cytoskeletal and mechanotransduction dynamics. For example, our data show that YAP nuclear translocation is observable within minutes of RhoA signaling activation. This is consistent with prior measurements from our group and others^15,31,50^ and coincides with the kinetics of cellular and cytoskeletal contractility. However, cytoplasmic translocation of YAP, after RhoA signaling termination, featured an 8 min delay followed by an efflux rate only half that of the influx rate. These data suggest that mechanoregulation of YAP nuclear efflux is governed by distinct mechanisms from nuclear influx and demand further study^50,51^. Similarly, opto-DLC1 enabled new insights into the kinetics and mechanisms of cytoskeletal remodeling. We observed rapid F-actin depolymerization of F-actin after RhoA termination. While multiple signaling nodes downstream of RhoA could play a role in decreased cytoskeletal tension, including ROCK-myosin contractility^52^ and mDia-profilin polymerization^53^, rapid depolymerization suggested a role for filament severing. Cofilins are F-actin-cleaving proteins that are phosphorylated by ROCK-LIM kinase signaling^54,55^ and dephosphorylated by Slingshot phosphatase (SSH)^56^. We found that SSH inhibition abrogated opto-DLC1-induced actin depolymerization. These data suggest that RhoA signaling inactivation unbridles cofilin from LIMK suppression. Optogenetic signaling termination therefore illuminates events that occur at time scales inaccessible to standard pharmacologic and genetic approaches.

Together, this toolbox broadens our ability to interrogate the spatiotemporal dynamics of GTPase signaling, particularly in processes and interactions in which inactivation or termination play a critical role, including cytoskeletal turnover-intensive cell migration^19^ and mutual-inhibitory crosstalk between RhoA and Rac1 signaling^14^. Future studies will use these tools to interrogate the spatial regulation of GTPase signaling termination, as we performed previously for RhoA activation.^31^

## METHODS

### Molecular cloning and protein engineering

To engineer optogenetic RhoA signaling termination tools, the catalytic GAP domains of DLC1/ARHGAP7^57^ and ARHGAP1^42^ identified using the PROSITE ExPASy database^58^ were human codon-optimized and synthesized as gBlocks™ by Integrated DNA Technologies (IDT). Domain arrangement combinations of BcLOV4, mCherry visualization tag, and the GAP domain were assembled as previously described^41^: BcLOV4 and mCherry were amplified from their mammalian codon-optimized reported fusion (Addgene plasmid #114595)^32^, and constructs were assembled by Gibson cloning NEB HiFi DNA Assembly Master Mix (E2621) into the pcDNA3.1 mammalian expression vector under the CMV promoter. Green tool variants were generated using a human codon-optimized eGFP gBlock™ synthesized by IDT. All genetic constructs were transformed into ultracompetent *E. coli* (New England Biolabs, C2984H). Rho biosensor dTomato-2xrGBD (plasmid #129625)^44^, miRFP703-tagged LifeAct (plasmid #79993)^59^, and EGFP-tagged YAP (plasmid #17843)^60^ were acquired from Addgene. Plasmid for BcLOV4-mCherry (plasmid #114595) and opto-GEF11-mCherry (plasmid #164473) are available through Addgene. Plasmids for BcLOV4-eGFP, opto-GEF11-eGFP, opto-DLC1, and opto-GAP1 will be available through Addgene.

### Cell culture and transfection

NIH 3T3 (ATCC, CRL1658) and HEK293T (ATCC, CRL3216) cells were cultured in D10 media composed of Dulbecco’s Modified Eagle Medium with Glutamax (Invitrogen, 10566016), supplemented with 10% heat-inactivated fetal bovine serum (FBS) and penicillin-streptomycin at 100 U/mL^−1^. Cells were maintained in a 5% CO2 water-jacketed incubator (Thermo/Forma Steri-Cycle, 370) at 37 °C. For imaging studies, cells were seeded onto poly-D-lysine-treated glass-bottom dishes (MatTek, P35GC-1.5-14-C) or gelatin-treated black 24-well glass bottom dishes (CellVis, P24-1.5H-N) at 25-30% confluency. Cells were transfected at ∼50–60% confluency 24 h later using the Lipofectamine 3000 transfection reagent for 3T3 (Thermo L3000008) or TransIT-293 (Mirus Bio, MIR-2700) for HEK293T, according to manufacturer instructions. For 3T3 cells, media was changed at transfection for antibiotic-free media, then replaced with full media ∼18 hours after transfection. Cells were imaged 24–72 h post-transfection.

### Optical hardware

Fluorescence microscopy was performed on an automated Leica DMI6000B fluorescence microscope under Leica MetaMorph or Micro-Manager control, with a sCMOS camera (pco.edge), an LED illuminator (Lumencor Spectra-X), and a 63× oil immersion objective. Excitation and illumination were filtered at the LED source (mCherry imaging, λ = 575/25 nm; eGFP imaging or wide-field BcLOV4 optogenetic stimulation, λ = 470/24 nm; miRFP703 imaging, λ = 632/22 nm). Fluorescent proteins were imaged with Chroma filters: mCherry (T585lpxr dichroic, ET630/75 nm emission filter, 0.2–0.5 s exposure), GFP (T495lpxr dichroic, ET 525/50 nm emission filter, 0.2 s exposure), and miRFP703 (AT655dc dichroic, ET655 nm emission, 0.5 s exposure).

### Tool construct screening and expression characteristics

For membrane recruitment quantification of BcLOV4 fusion tool constructs, prenylated GFP was co-transfected as a membrane marker. An mCherry fluorescence image (500 ms exposure) was captured to assess protein expression level and subcellular distribution. Cells were then illuminated with a 5 s-long blue light pulse to stimulate BcLOV4 membrane recruitment, during which the GFP membrane marker was imaged. Cells were then imaged immediately following blue light stimulation. Membrane localization and dissociation were measured by line section analysis and correlation with prenylated GFP in Fiji and Python.

### Quantification of RhoA signaling kinetics

For association studies, 3T3 cells in 35 mm confocal dishes were transfected with plasmids encoding dTomato-2xrGBD biosensor and either eGFP-tagged BcLOV4 control, opto-GEF11, or opto-DLC1 in a 1:1 ratio for a total DNA mass of 2500 ng per dish. One field of view per dish was imaged for 60 seconds, alternating between a 200 msec eGFP imaging/BcLOV4 stimulation scan and a 400 msec dTomato imaging scan. For dissociation studies, cells were transfected with 1250 ng of either dTomato-2xrGBD biosensor or mCherry-tagged BcLOV4 control, opto-GEF11, or opto-DLC1. One minute of stimulation was performed as in association studies, followed by a 5-minute imaging window in which a 400-msec mCherry/dTomato scan was performed every 5 seconds. Membrane and peri-membrane cytosol regions were isolated via image binarization and object subtraction in Fiji and per-region average pixel intensities were measured at each imaging timepoint.

### Quantification of actin polymerization and depolymerization

For actin time course imaging, cells were transfected with a 1:1 ratio of mCherry-tagged BcLOV4 control, opto-GEF11, or opto-DLC1 and miRFP703-tagged LifeAct. Each field of view was imaged for 30 minutes, with cells stimulated with a 500 msec blue-light pulse every 15 seconds. mCherry (tool) and miRFP (LifeAct) were each imaged once per minute with 500 msec exposure. Time course images were bleach-corrected by exponential fit in Fiji and mean LifeAct-miRFP intensity for each cell was measured every minute. For inhibitor studies, cells were treated with slingshot inhibitor D3 (Bepharm Scientific, A1516343, 5 µM in DMSO), or DMSO control for 30 minutes prior to imaging. For opto-GAP1 actin visualization, cells were washed with PBS and the media was replaced with DMEM supplemented with penicillin-streptomycin and without FBS 24 hours after transfection. Light-exposed plates were incubated under Arduino-controlled blue strip LEDs (light intensity 15 mW cm^−2^) strobing at a 1.6% stimulation duty cycle in a 5% CO_2_ water-jacketed incubator for four hours. Dark-adapted controlled cells were incubated in the same incubator in a foil-wrapped plate to prevent light exposure. Cells were fixed with 4% paraformaldehyde in PBS at room temperature for 10 min, washed twice with PBS, and then permeabilized with 0.1% Triton-x-100 in PBS for 15 min. Cells were blocked with 1% BSA in PBS for 30 min, then stained with Alexa Fluor 488 Phalloidin (Invitrogen, A12379) diluted 1:400 in PBS. Plates were washed twice prior to imaging. Filamentous actin level was quantified by normalizing total-cell Alexa Fluor 488 fluorescence to cell area.

### Quantification of YAP translocation kinetics

Cells were transfected with a 1:1 ratio of mCherry-tagged BcLOV4 control, opto-GEF11, or opto-DLC1 and eGFP-tagged YAP. mCherry (tool) and eGFP (YAP; doubles as blue light tool stimulation) were each imaged every 15 seconds with 500 msec exposure. Time course images were bleach-corrected by exponential fit in Fiji and mean YAP nuclear and cytosolic intensities for each cell were measured every minute.

### Statistical analysis

Statistical analysis was conducted in GraphPad Prism version 10.5.0. One-way ANOVA with multiple comparison tests (Tukey or Šidák) or two-tailed Student’s t-tests were used to determine statistical significance, with a P-value of less than 0.05 considered significant. Sample sizes are indicated in figure legends and graphs display individual data points as scatterplots. Kinetic constants were calculated using one-phase association and dissociation curve fitting.

## ACKNOWLEDGEMENTS

The authors would like to thank all members of the Boerckel Lab for constructive discussions and the Perelman School of Medicine Cell and Developmental Biology Microscopy Core (RRID SCR_022373) for confocal microscope access.

## FUNDING

National Science Foundation Louis Stokes Alliances for Minority Participation (LSAMP) (J.B.)

National Institutes of Health grant P30 AR069619 (J.D.B.)

National Institutes of Health grant R01 GM143400 (J.D.B.) including a Research Supplement to Promote Diversity in Health-Related Research (P.C.S.)

Penn Achilles Tendinopathy Center of Research Translation P50 AR080581 (J.D.B.)

## AUTHOR CONTRIBUTIONS

E.E.B. and J.D.B. conceived and supervised the research and designed the experiments. E.E.B, J.B., and P.C.S. performed experiments and analysis. E.E.B and J.D.B. wrote the manuscript. All authors discussed and revised the manuscript.

## Supplementary Figures

**Supplementary Figure 1.**
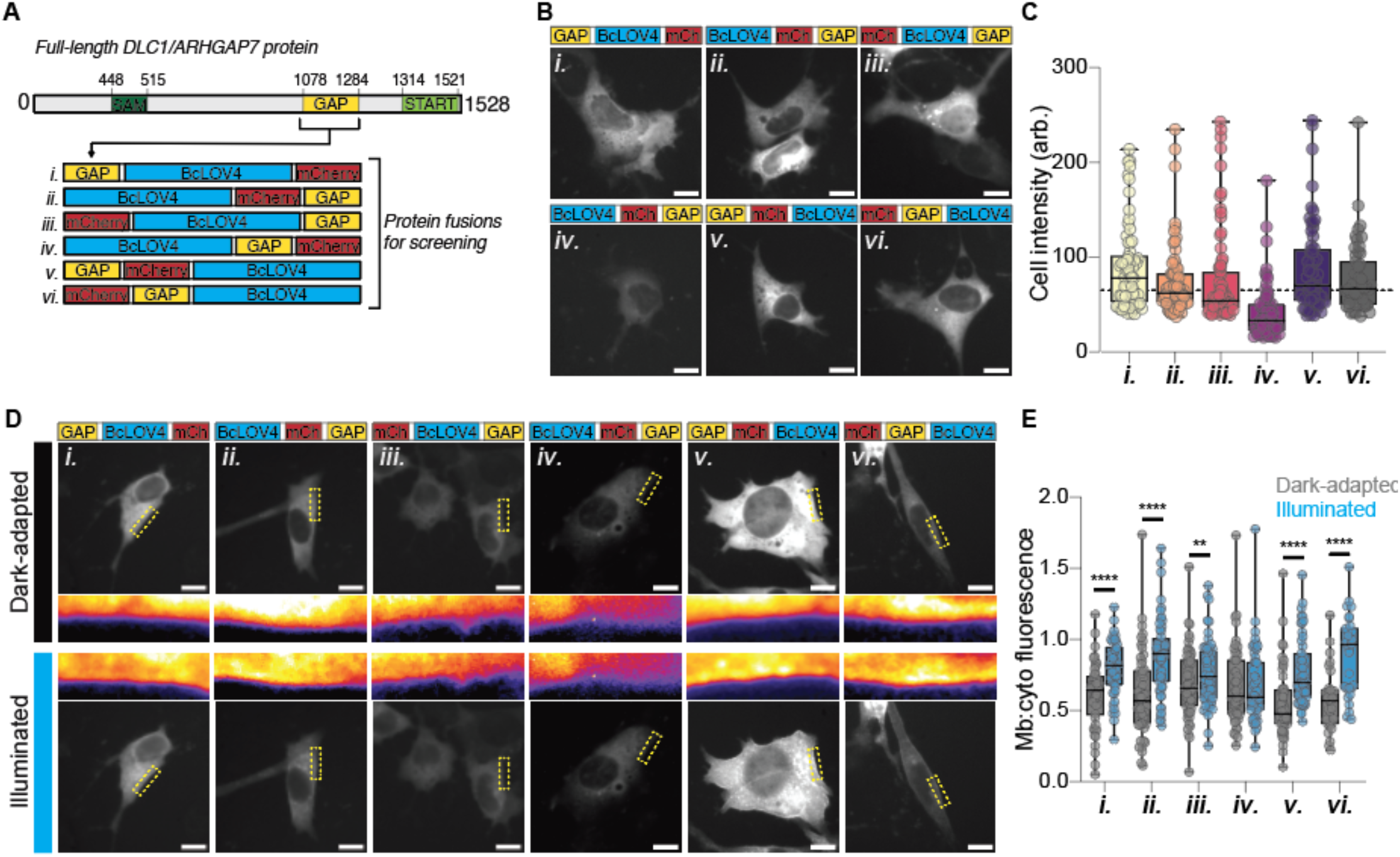
Engineering optogenetic DLC1. **A**) Schematic of catalytic GAP domain of deleted in liver cancer 1 (DLC1/ARHGAP7) protein used in optogenetic protein fusions. SAM = sterile alpha motif; START = StAR-related lipid transfer; GAP = GTPase accelerating protein. BcLOV4, mCherry, and DLC1-GAP domains were separated by flexible (GGGS)_2_ linkers. **B**) Fluorescence micrographs showing representative expression patterns of domain arrangements in 3T3 cells in the dark-adapted state. Scale = 10 µm. **C**) Expression level measured by mCherry fluorescence of genetic constructs. Dashed line = mean BcLOV4-mCherry fluorescence intensity. N = 53– 81 cells per condition. **D**) Representative membrane localization of DLC1 protein fusions before and after blue light stimulation. Inset shows boxed region in perceptually uniform color palette. **E**). Ratio of membrane-localized vs. cytosolic protein for each screened protein in dark-adapted and blue light-illuminated states. N = 30–60 cells per condition. Paired t-test: (**) p < 0.01, (****) p < 0.0001.

**Supplementary Figure 2.**
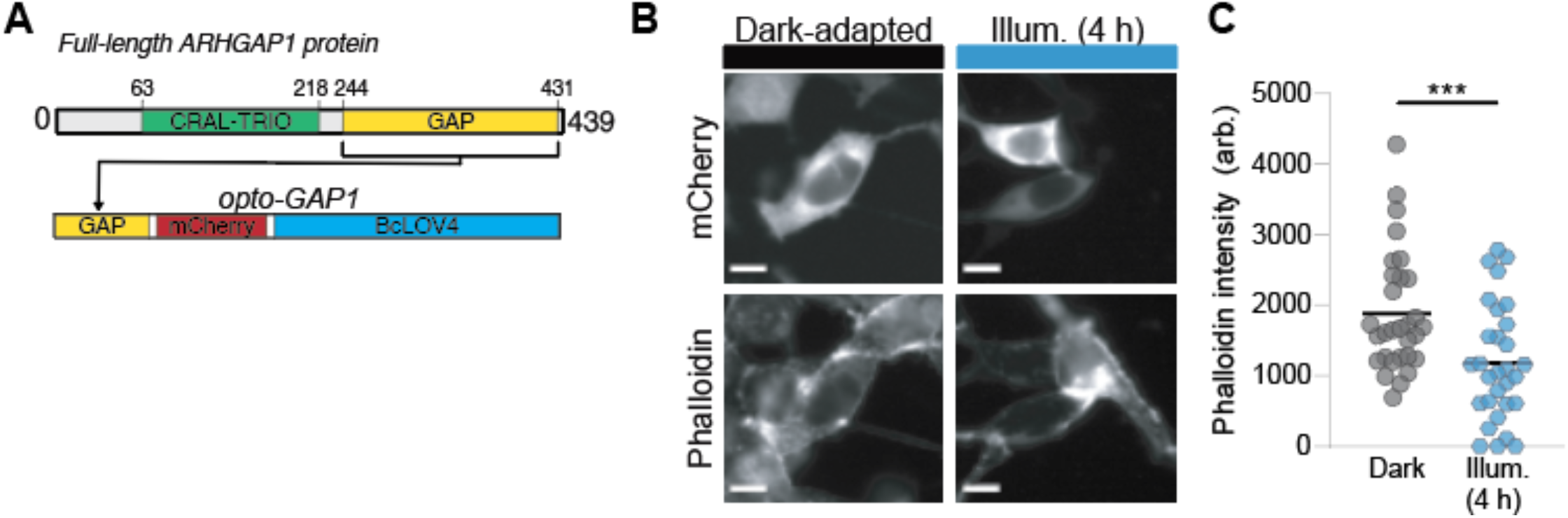
Engineering opto-GAP1 using ARHGAP1. **A**) Schematic of catalytic GAP domain of ARHGAP1 protein used in optogenetic tool. CRAL-TRIO = lipid binding domain; GAP = GTPase accelerating protein. BcLOV4, mCherry, and ARHGAP1-GAP domains were separated by flexible (GGGS)_2_ linkers to create opto-GAP1. B. HEK293T cells transiently transfected with opto-GAP1 were fixed following four hours of incubation either in the dark or under pulsatile blue light stimulation. mCherry (top) shows tool localization in transfected cells; phalloidin (bottom) shows F-actin in all cells. Scale = 10 µm. C. Quantification of average phalloidin intensity per cell. N = 30 cells per condition. (***) p < 0.001, Mann-Whitney U test.

**Supplementary Figure 3.**
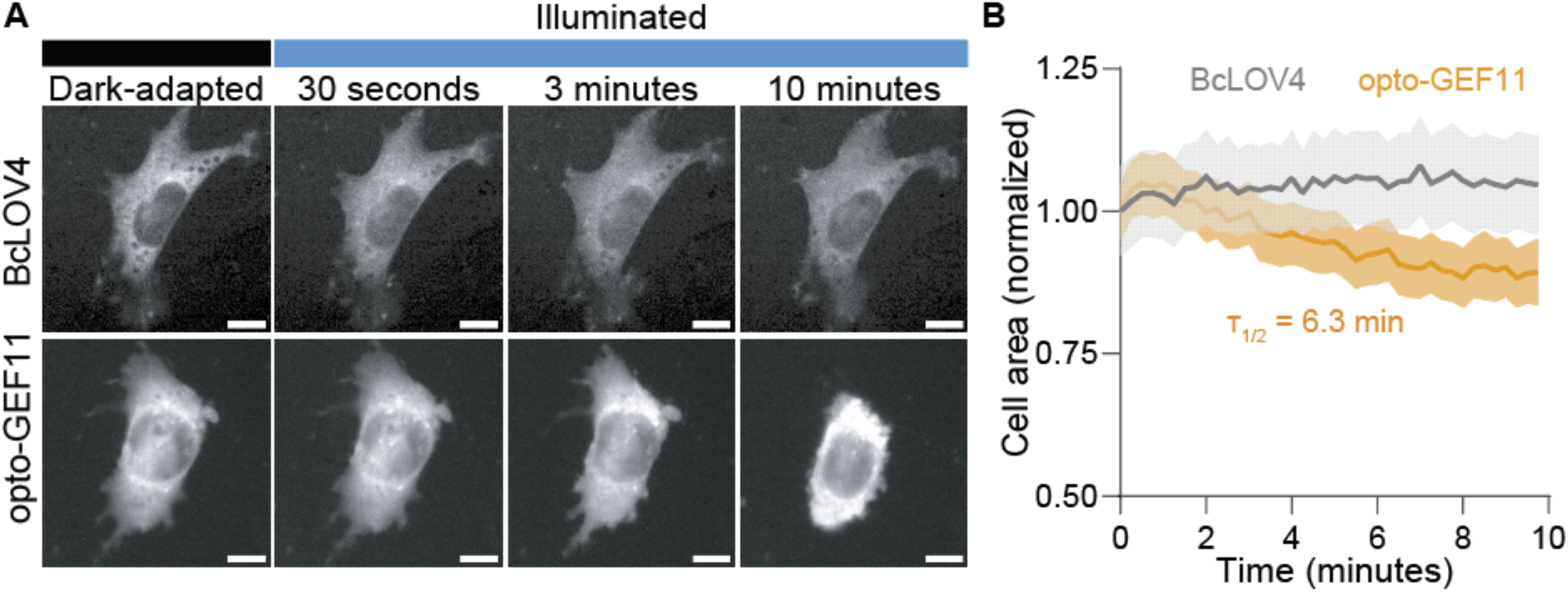
Demonstration of opto-GEF11-mediated cell contraction in 3T3 fibroblasts. **A**) Epifluorescence micrographs of 3T3 fibroblasts expressing either BcLOV4 (top) or opto-GEF11 (bottom), visualized by fused mCherry tag, before and during 10 minutes of pulsatile blue light stimulation (1.67% duty ratio). Scale = 10 µm. **B**) Quantification of relative area of stimulated cells, normalized to cell area prior to illumination. Mean ± S.E.M, N = 40 cells per condition. Time constant calculated by one-phase dissociation exponential fit.

**Supplementary Figure 4.**
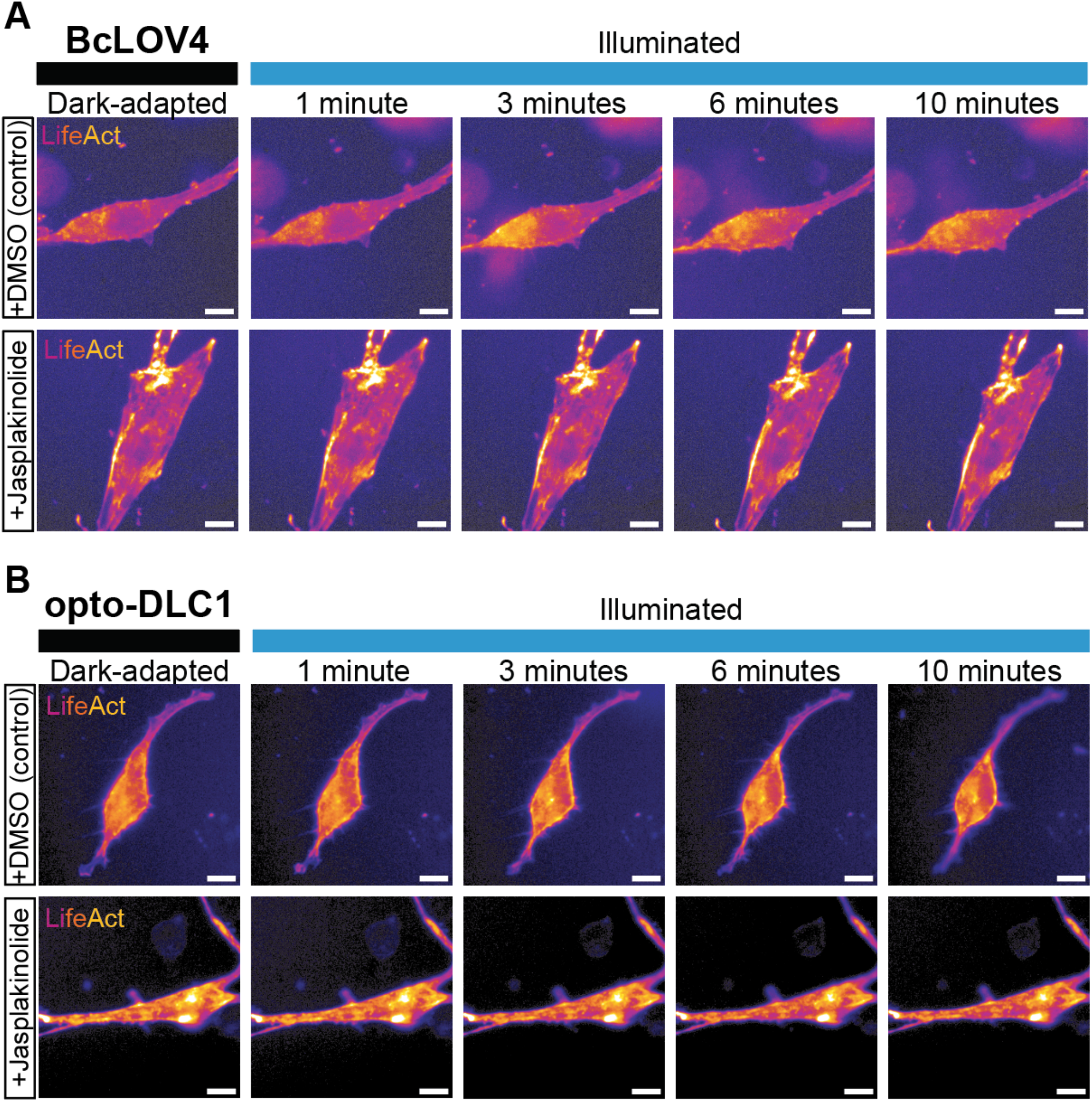
Representative images of opto-GEF11. (**A**) and opto-DLC1 (**B**) co-expressed with LifeAct-miRFP703 during a 10-minute period of pulsatile blue light stimulation, following 30 minutes of incubation with (top) DMSO or (bottom) jasplakinolide. Scale = 10 µm.

**Supplementary Table 1.** Time constants and plateau values.

| Figure | Condition | Parameter | Value | 95% confidence interval |
| --- | --- | --- | --- | --- |
| 1 | BcLOV4 | Tool association $\tau_{1/2}$ | 0.7 s | 0.3–1.2 s |
| 1 | opto-GEF11 | Tool association $\tau_{1/2}$ | 1.0 s | 0.4–2.1 s |
| 1 | opto-GEF11 | Rho biosensor $\tau_{1/2}$ | 8.4 s | 1.3–nd s |
| 1 | opto-DLC1 | Tool association $\tau_{1/2}$ | 1.7 s | 1.2–2.5 s |
| 1 | opto-DLC1 | Rho biosensor $\tau_{1/2}$ | 1.8 s | 0.9–4.0 s |
| 2 | BcLOV4 | Tool dissociation delay | 19.1 s | 16.6–22.6 s |
| 2 | BcLOV4 | Tool dissociation $\tau_{1/2}$ | 5.3 s | 5.2–11.1 s |
| 2 | opto-GEF11 | Tool dissociation delay | 19.3 s | 15.5–24.5 s |
| 2 | opto-GEF11 | Tool dissociation $\tau_{1/2}$ | 5.9 s | 5.3–13.2 s |
| 2 | opto-GEF11 | Rho biosensor delay | 37.0 s | nd |
| 2 | opto-GEF11 | Rho biosensor $\tau_{1/2}$ | 37.0 s | 1.1–212.6 s |
| 2 | opto-DLC1 | Tool dissociation delay | 17.4 s | 13.3–19.4 s |
| 2 | opto-DLC1 | Tool dissociation $\tau_{1/2}$ | 7.7 s | 8.1–15.8 s |
| 2 | opto-DLC1 | Rho biosensor delay | 42.1 s | 21.3–nd s |
| 2 | opto-DLC1 | Rho biosensor $\tau_{1/2}$ | 7.6 s | 1.6–19.4 s |
| 4 | opto-GEF11 | YAP N:C ratio $\tau_{1/2}$ | 12.6 s | 6.5–80.3 s |
| 4 | opto-DLC1 | YAP N:C ratio delay | 8.6 s | 1.9–13.5 s |
| 4 | opto-DLC1 | YAP N:C ratio $\tau_{1/2}$ | 22.5 s | 2.0–nd s |
| 5 | opto-GEF11 | F-actin $\tau_{1/2}$ | 2.0 s | 0.3–nd s |
| 5 | opto-DLC1 | F-actin $\tau_{1/2}$ | 3.4 s | 0.6–15.7 s |
| S2 | opto-GEF11 | Area change $\tau_{1/2}$ | 6.3 s | 1.9–nd s |

